# Two new ant species of the genus *Camponotus* (Hymenoptera: Formicidae) from the Andean Region of Chile

**DOI:** 10.1101/2025.08.26.672482

**Authors:** Benjamín Arenas-Gutierrez, Carlos P. Muñoz-Ramírez

## Abstract

*Camponotus* (Hymenoptera: Formicidae) is one of the most diverse ant genera in the world (>1000 species); however, only six species have been described in Chile, with the last description being in 1935. Recently, various specimens have been collected that do not match the descriptions of known species, suggesting the existence of new species that need to be described. In the present study, two new species are proposed for Chile. Descriptions, diagnoses, and photographs are included for the new taxa. With these new species, the myrmecological fauna of the genus *Camponotus* present in the country is increased by 33%.

## Introduction

The genus *Camponotus* (Hymenoptera, Formicidae) belongs to the tribe Camponotini, in the subfamily Formicinae. It is one of the most diverse ant genera in the world, with more than 1,000 species described to date (Antweb, 2024), and is present in all regions of the world (except in the polar regions), especially in the Neotropical regions and North America (Bolton, 1995). Morphologically, they are large ants (between 5 and 25 mm) characterized by having, as with the entire subfamily, only a single petiolar node (petiole) in the form of a scale and a structure called an acidopore on the seventh abdominal sternite, which is used to expel formic acid when threatened. Specifically, members of the genus *Camponotus* display an acidopore without a ring of hair, lack a metapleural gland, and exhibit intraspecific polymorphism, with major workers possessing large heads that can exceed the size of minor workers by two to four times (Fisher & Cover, 2007). The taxonomy of this group is recognized for its complexity due to the large number of species and their high morphological variation both within and between species, which has made it difficult to delimit species and to understand their phylogenetic relationships (Mackay and Delsinne, 2009).

In Chile, the genus *Camponotus* comprises six species described to date, a low number compared to the number of species in neighboring countries (Bezděčková et al., 2015; Kusnezov, 1951). The first *Camponotus* species described for Chile are those documented by Spinola in the classic text by Gay, “Historia Física y Política de Chile,” volume 6 (1851): *Camponotus chilensis*, *C. distinguendus*, and *C. ovaticeps*, which at that time were classified under the genus *Formica*. Seven years later, Smith (1858) described *Camponotus morosus* (as *Formica morosa*). Subsequently, Roger (1863) described a fifth species native to the Chillán mountain range, which was named in honor of Spinola as *Camponotus spinolae*. Finally, Menozzi (1935) described the most recently known species to date, *Camponotus hellmichi*, originally described as a variety of *Camponotus morosus*.

In a recent field exploration of the Andean area of the Maule Region (figure 1), the first author collected several specimens of *Camponotus* exhibiting some characteristics that did not fit the available descriptions for the Chilean species. Subsequently, a detailed examination concluded that these specimens belong to two undescribed species. In this article, we name the two new species and provide detailed descriptions and diagnosis, including photographs of all available castes.

**Figure 1:**
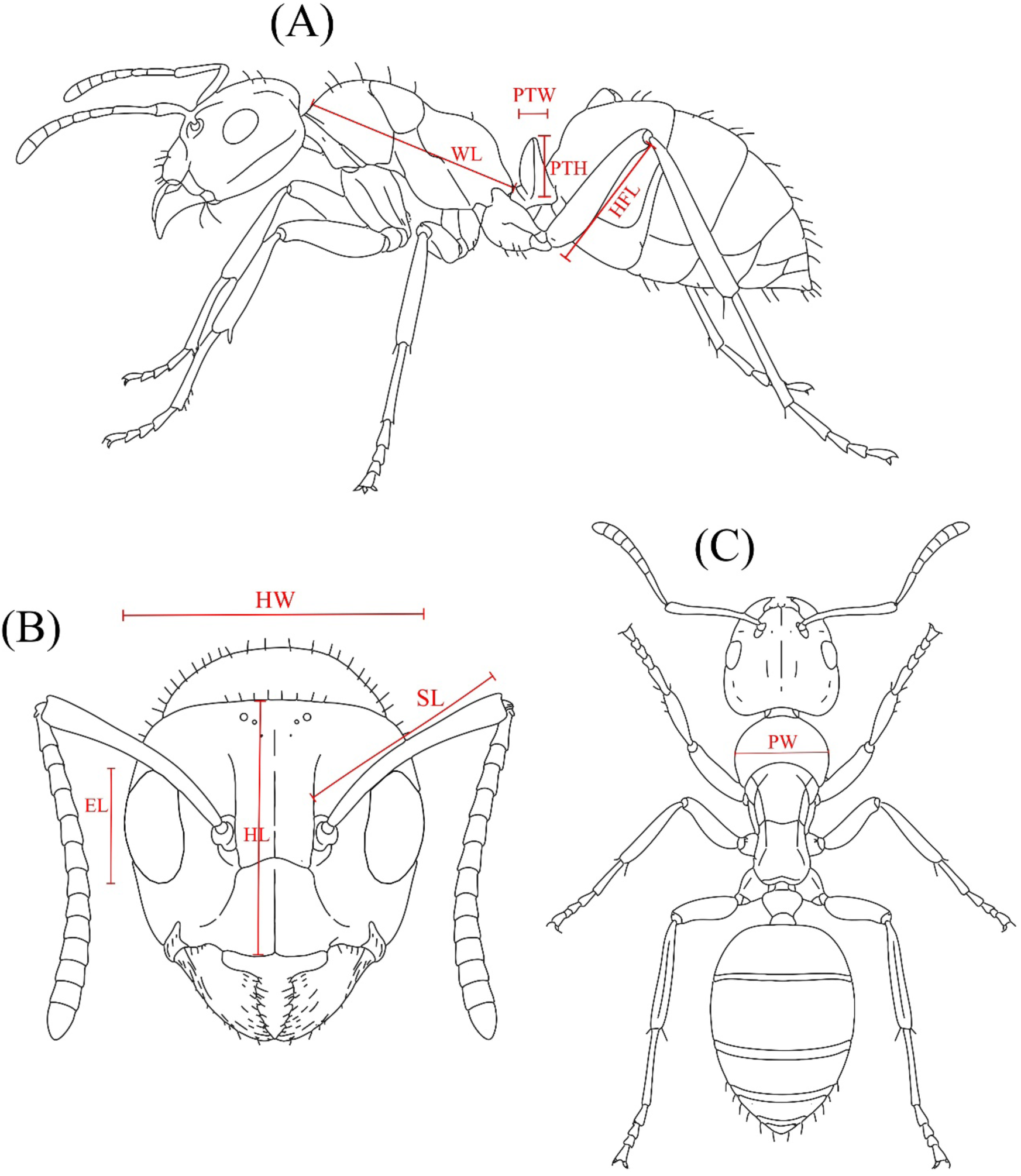
Scheme of a general ant depicting measurements on specimens: (A) lateral view, (B) head in frontal view, and (C) dorsal view. HL: Head Lenght; HW: Head width; SL: Scape Lenght; EL: Eye lenght; PW: Pronotum Width; PTH: Petiole Height; PTW: Petiole Width; WL: Weber’s lenght; HFL: Hind Femur Lenght.

**Figure 2:**
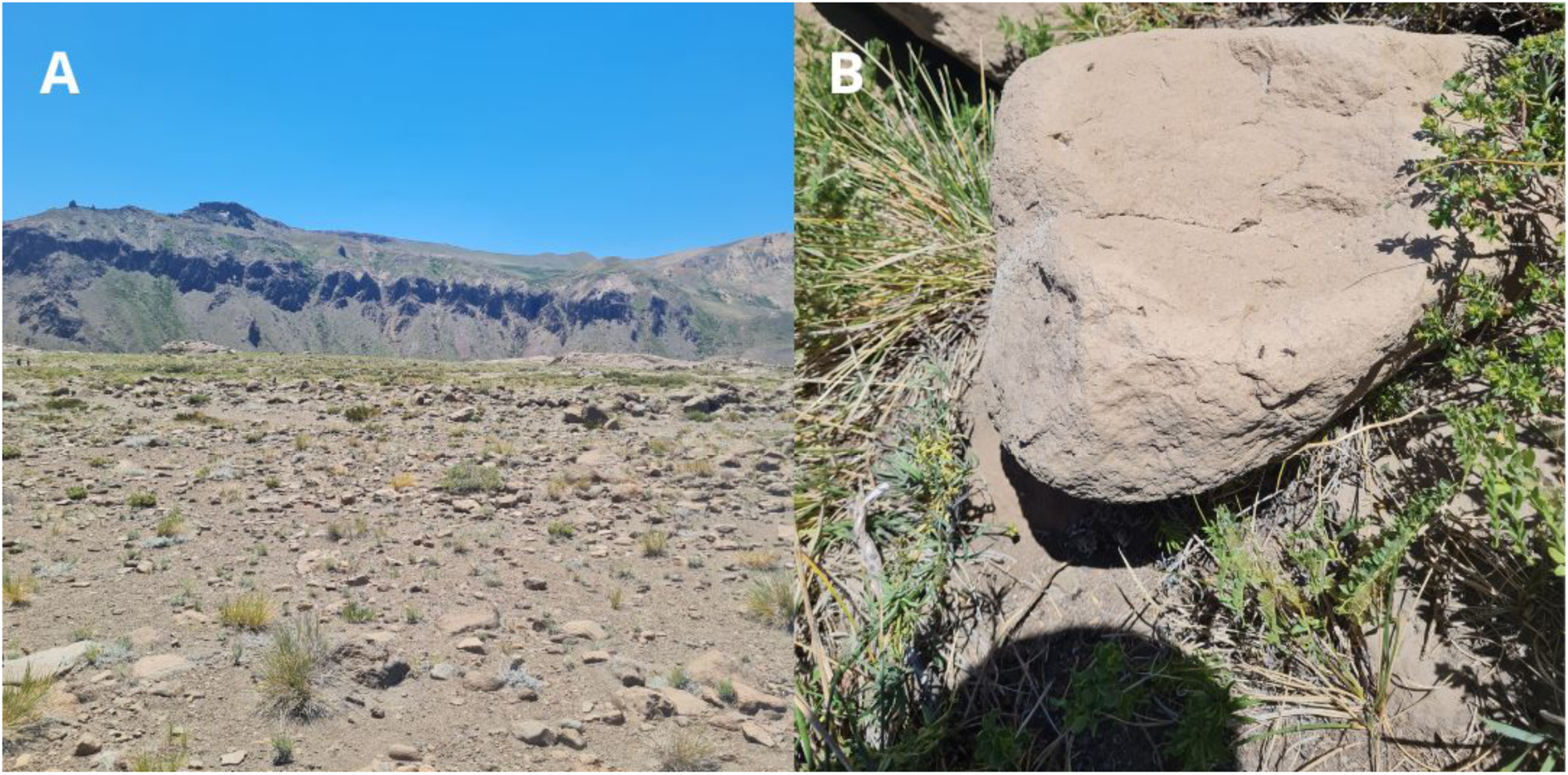
Habitat of *Camponotus carminatus* sp. nov. and *Camponotus brunnipes* sp. nov. from the locality of “Maule Alto” (Cascada Invertida, 1850 masl). (A) General view of the Andean steppe-type ecosystem where the samples were collected; (B) Rock under which one of the *C. carminatus* sp. nov. nest was found. In the photograph, *C. carminatus* sp. nov. individuals can be seen wandering around the rock.

## Material and methods

Specimens were collected at the “Cascada Invertida” site (San Clemente, Maule Region, 1800 m above sea level) on two occasions, February 2024 and April 2025. The explored habitat was mainly high Andean steppe (figure 1b), where the main collection method was manual gathering, complemented by captures via a mouth aspirator. Individuals were collected from beneath the rocks where their nests were located. All specimens were preserved in 99% ethyl alcohol (EtOH), ensuring that only individuals from the same colony were stored in each container. After this, they were transported and stored at −20 °C in the Insect Molecular Ecology Laboratory (LEMIn) of the Instituto de Entomología, Universidad Metropolitana de Ciencias de la Educación (UMCE-IE). Sampling in 2024 was carried out by Benjamín Arenas and Pablo Arenas, while sampling in 2025 was carried out by Benjamín Arenas and Isabel Sánchez. The geographic coordinates indicate the general collection site, not the exact collection points. The collections were made within a 100-meter radius of the indicated geographic point.

### Institutional abreviations

**UMCE-IE:** Instituto de Entomología, Universidad Metropolitana de Ciencias de la educación, Santiago, Chile.

**MNHN:** Museo Nacional de Historia Natural, Santiago, Chile.

**UNAP:** Universidad Arturo Prat, Iquique, Chile

### Measurements

The specimens were photographed using a Celestron digital microscope, model “Micro Direct.” Photographs of all castes (as available) were taken for each species in lateral, cephalic, and dorsal views. A piece of graph paper was included when taking photographs to serve as a reference for subsequent measurements. In each photograph, a white line representing 1 mm was included as a scale.

Morphological measurements were conducted using the Tpsutil32 and Tpsdig264 programs (Rohlf 2010), calibrating each photograph with the graph paper included in each photo. All measurements are presented in millimeters (mm), first indicating their arithmetic mean, followed by the minimum and maximum values within parentheses.

### Measurement Abbreviations

**HL**: “Head Length”, measured in frontal view; straight line from the midpoint of the anterior border of the clypeus to the midpoint of the posterior border of the vertex.

**HW**: “Head Width”, measured in frontal view; straight line from one lateral edge to the other, passing just posterior to the eyes.

**SL**: “Scape Length”, measured in frontal view; straight line from the joint with the funiculus to the base of the scape, excluding the condylar bulb.

**EL**: “Eye Length”, measured in frontal view; maximum diameter of the compound eye.

**PW**: “Pronotum Width”, measured in dorsal view, maximum width of the pronotum.

**PTH**: “Petiole Height”, measured in lateral view; straight line, transverse to the ant’s body, from the edge of the sternal plate (base of the petiole) to the apex of the petiole.

**PTW**: “Petiole Width”, measured in lateral view; maximum width of the petiolar node in lateral view.

**WL**: “Weber’s length” or “Mesosomal length”, measured in lateral view; measured as a diagonal line from the edge where the pronotum meets the cervical shield to the posterior basal angle of the metapleuron.

**HFL**: “Hind femur length”, measured in dorsal view; straight line from the trochanter-femur joint to the femur-tibia joint.

### Index abreviations

**CI:** “Cephalic index”; HL/HW

**SI:** “Scape index”; SL/HL

**PI:** “Petiole index”; PTH/PTW

**FI:** “Femur index”; HFL/WL

To refer to the inclination of the pilosity, the terms used in Wilson (1955) were employed. “Appressed”: 0-5°; Decumbent: 5-20°; Sub-decumbent: 20-60°; suberect: 60-80°; erect: 80-90°. All degrees are relative to the tegument surface.

The article by Salata et al. (2023) was followed for the morphological description of the new species, taking into consideration the guidelines suggested by Braby et al. (2024) for the description of new species in zoology.

## Results

### 1) Camponotus carminatus sp nov

(Figs. 3-5)

**Fig. 3:**
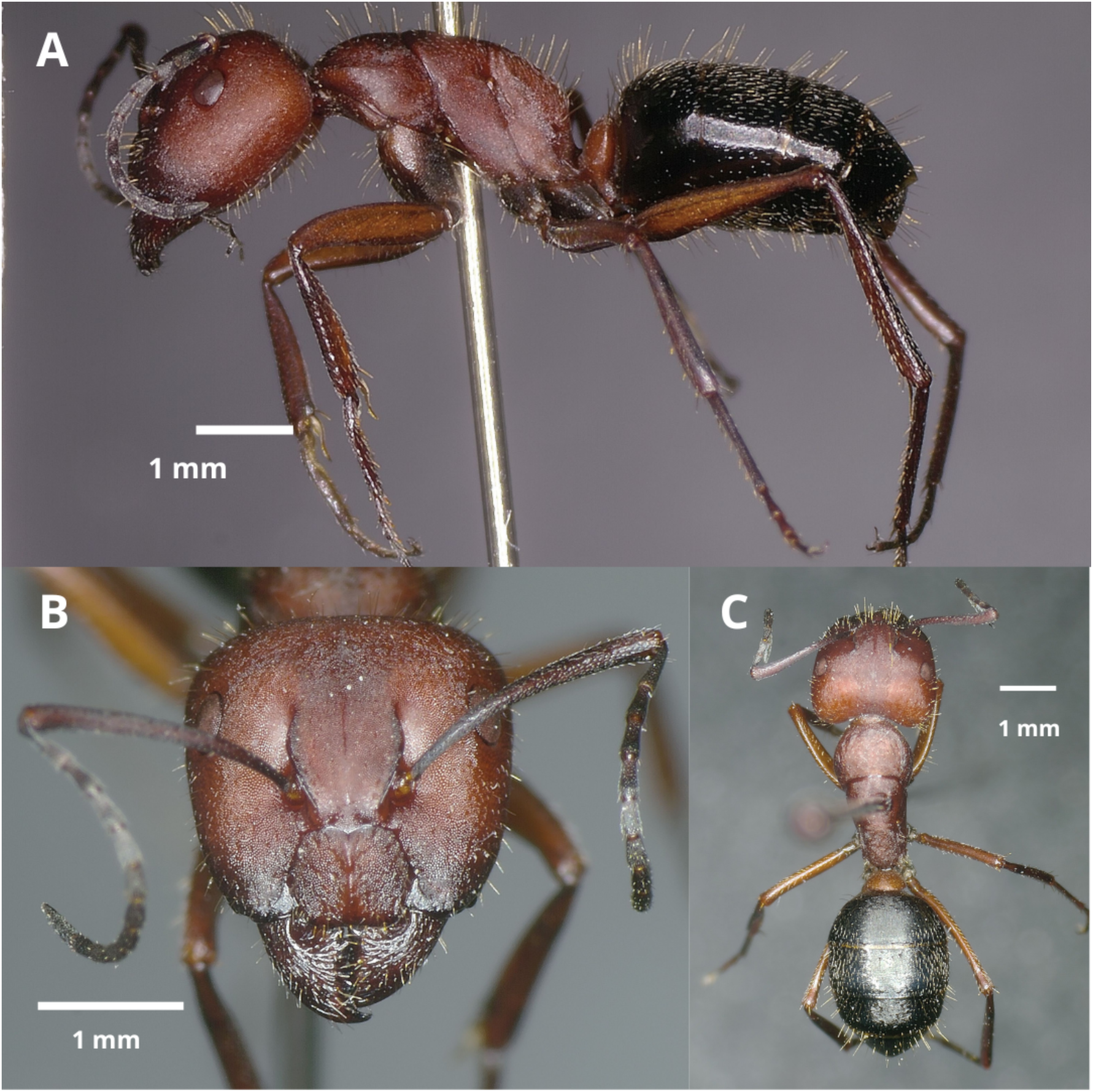
C*a*mponotus *carminatus* sp nov. major worker photographies (A) Lateral view (B) Frontal / Cephalic view (C) Dorsal view

**Fig. 4:**
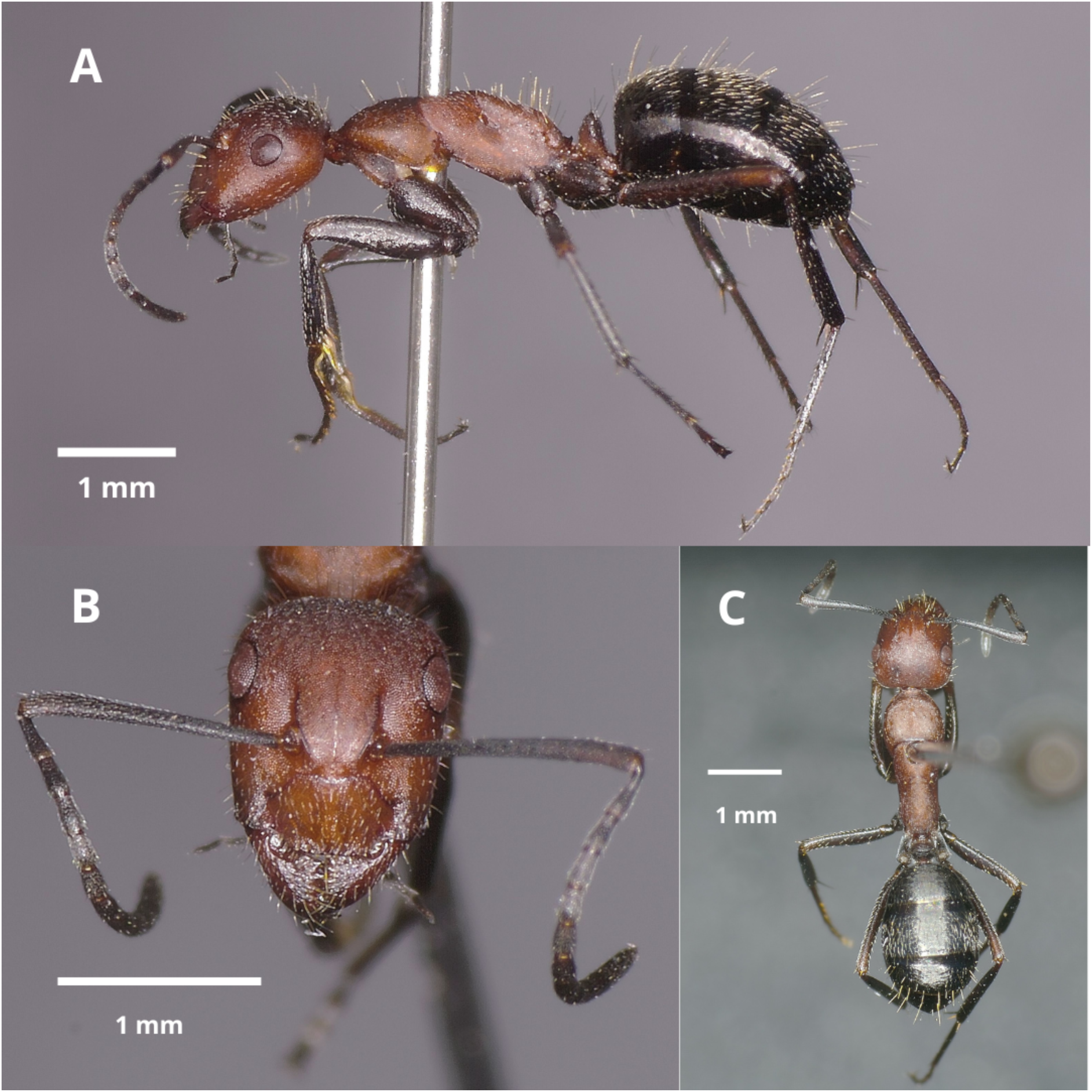
C*a*mponotus *carminatus* sp nov. minor worker photographies (A) Lateral view (B) Frontal / Cephalic view (C) Dorsal view

**Fig. 5:**
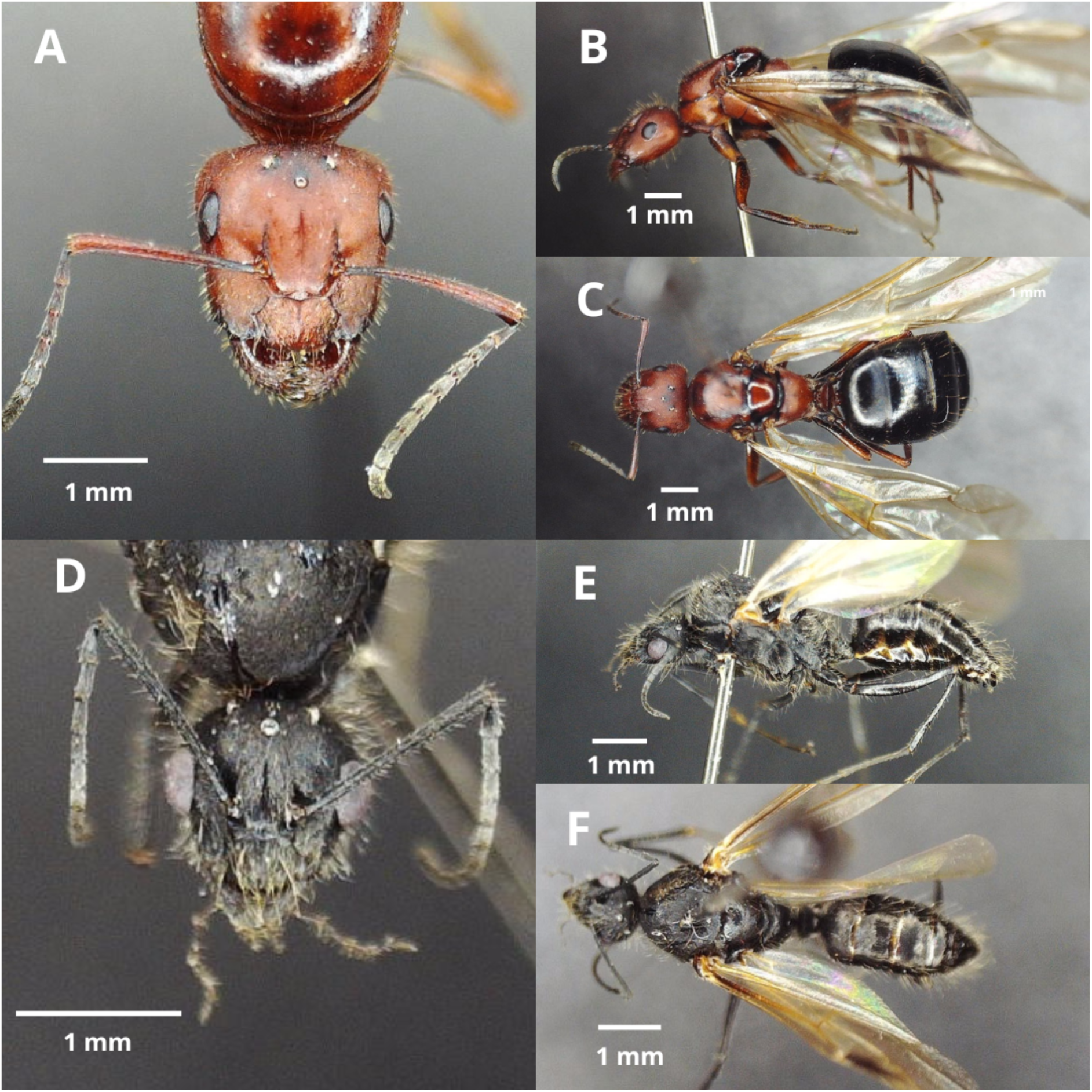
C*a*mponotus *carminatus* sp nov. winged female and male photographies (A) Winged female frontal / cephalic view, (B) Winged female lateral view, (C) Winged female dorsal view, (D) Male Frontal / Cephalic, (E) Male lateral view, (F) Male dorsal view.

**Type material (pinned): Holotype:** Major worker (UMCE_IE_1130). Chile, San Clemente (Maule Region), Cascada Invertida | −35.95473 / −70.57363, 10 IV 2025, B. Arenas (UMCE-IE, UNAP). 3 Paratypes (UMCE_IE_1131 - UMCE_IE_1133): 2 major workers, 1 minor worker, same data as holotype (IE-UMCE). 5 paratypes (UMCE_IE_1134 - UMCE_IE_1138): 3 minor workers, 1 winged female, 1 male: Chile, San Clemente (Maule Region), Cascada Invertida | −35.95473 / −70.57363, 28 II 2024, B. Arenas (UMCE-IE, UNAP).

**Etimology:** The epithet “carminatus” refers to the intense red color, similar to carmine, that individuals of this species exhibit on their head and mesosoma.

**Diagnosis:** *Camponotus carminatus* is a species whose workers are characterized and distinguished from the workers of all other species present in Chile by possessing the following group of traits: head and thorax with reddish coloration, a feature present in all its castes except the males. Scape and antennal funiculus with reddish coloration. It has pilosity at the edges of the gena (setae). Legs with black coloration and reddish areas, mainly close to the joint regions. The gaster has sparse pilosity (setae) on the tergal surface, which allows the visibility of the black-colored integument.

#### Description

**Major worker** (n=5, Figs. x-x): **Measurements:** EL: 0.39 (0.36-0.43); HL: 1.994(1.76-2.22); HW: 2.12 (1.94-2.27); SL: 1.798 (1.62-1.96); PW: 1.31 (1.22-1.43); PTH: 0.768 (0.7-0.88); PTW: 0.616 (0.51-0.73); WL: 2.754 (2.6-2.9); HFL: 1.938 (1.81-2.05); CI: 0.9389 (0.9072-0.9779); SI: 0.9037 (0.8468-0.9378); PI: 1.0951 (0.8082-1.2464); FI: 0.7036 (0.6931-0.7269)

**Color**: Head almost entirely red, slightly opaque, with darkening on the lower portion of the gena, the area adjacent to the anterior mandibular joint, the anterior margin of the clypeus, and along the entire frontal carina. The mesosoma and petiolar scale have the same red color as the head. Gaster predominantly black, with slightly yellowish hues on the edges of the sternal and tergal plates. Mandibles and antennae are a darker red than the rest of the head, especially on the antennal funiculus. Legs have black lines along their entire length, which are most pronounced on the coxa.

**Head**: Slightly wider than tall, subquadrate, with subparallel sides that are broader behind the eyes and the gena converging forward; occipital edges rounded, forming an angle of nearly 90 degrees with the lateral edge. Anterior and posterior margins of the clypeus bilobed, with a notch in the central sector; in the medial sector of the anterior margin of each lobe there are 3 to 5 yellowish setae, and in the lateral portion a similar number but smaller (2 to 3 times smaller) and of a more whitish coloration; the tegument of the clipeal plate is micropunctate and with little shine, with small and subtle concavities scattered sparsely over its surface; presence of both long and short setae (the latter 3 to 4 times shorter), the first ones erect and the second ones pressed against the surface (“apressed”) to the point that their visibility is difficult; presence of a medial keel, which reaches the anterior margin of the clypeus, extending toward the posterior margin where it usually bifurcates towards each lobe. Eyes relatively small compared to the size of the head, and relatively oval, approximately twice as tall as they are wide, and 0.45 times the height of the gena. Frontal carina extending from the upper sector at the margin of the clypeus to about halfway up the eyes, slightly arched; frontal plate with a thin and superficial central groove, which extends towards the coronal sector just above eye level, where it is accompanied by a slight depression and darkening of the tegument; tegument surface similar to the clypeus, likewise with both long and short setae. Antennal fossa shallow, without special sculpturing. Head with tegument similar to that of the clypeus, with long-erect and short-“apressed” setae over its whole surface, including the lateral and posterior edges. Antennal scapes moderately long; with visible anteroposterior curvature in dorsal view, approximately 0.85–0.94 times the height of the head (SI), with its apex gradually widening; funiculus longer than the scape; pedicel slightly elongated with respect to the first flagellar segment. Scape with very fine micro-reticulation, with lines so fine as to be difficult to perceive, tegument matte; with small notches sparsely scattered over its surface, similar to the head’s tegument; presence of short whitish appressed pilosity; in frontal view, the lower sector of the antennal condylar bulb with abundant, very small erect setae, approximately 10 times smaller than the erect setae of the clypeus. Mandibles rounded, with micro-reticulations increasing towards the distal area, becoming very thick in the area adjacent to the teeth; tegument with small spaced notches and setae of intermediate size compared to the erect setae present on the clypeus, subdecumbent in the direction of the mandibular apex, and small “apressed” setae scattered over the entire surface. **Mesosoma**: Long, 2.03 - 2.15 times longer than wide; dorsum of mesosoma forming a regular arch between pronotum, mesonotum, and propodeum, without any indentation between them; propodeum without protrusions. Pronotum rounded at its lateral edges; anterior margin flexed towards the dorsal. Mesosoma densely micropunctate and slightly opaque; anterior margin of pronotum (neck) microstriated; propodeum micropunctate and microstriated. Numerous erect setae on the dorsal area of the pronotum, mesonotum, and propodeum, relatively more abundant in the latter but not on the sides; “appressed” setae over the entire surface of the pronotum, mesonotum, and propodeum, both dorsally and laterally. **Petiole**: Scale-shaped, thin, 1.93 - 2.3 times thinner than wide; apex rounded along its entire length; posterior surface flat, anterior surface slightly convex; presence of 8 long setae, similar in size to those on the clypeus, on the edge of the petiolar scale, separated into two groups of four by a slight bare recess in the medial area; surface displays faintly opaque microreticulation. **Gaster**: Tegument with very fine micro-reticulation, surface slightly opaque except for the edges of tergal plates, which are shinier; Presence of long, erect setae and short, appressed setae across the entire dorsal and ventral surface, distributed sparsely to allow the tegument to be visible; presence of a line of setae of intermediate size (¾ to ½ the length of the long setae) situated along the entire posterior edge, close to each plate, both on tergal and sternal plates. **Legs**: Posterior femur elongated, 0.69– 0.73 times as long as the mesosoma (FI); femur of all legs slightly flattened anteroposteriorly, tibia also flattened but to a lesser extent; coxa, femur, and tibia covered with long erect setae, reaching the same size on the coxa and ⅓ or ½ the length of the clypeal setae on the femur and tibia; presence of short pilosity, appressed to the integument, along the entire length of the legs; femur, in the area of the femoro-tibial articulation, with four setae: one anterior, one posterior, and two on the dorsal side of the joint, separated from each other—present on all legs. Integument micropunctate and microstriated along the entire length of the legs.

**Minor worker** (n=5, Figs. z-z): **Measurements:** EL: 0.302 (0.28-0.32); HL: 1.28(1.22-1.34); HW: 1.16 (1.08-1.23); SL: 1.304 (1.24-1.4); PW: 0.902 (0.83-1); PTH: 0.566 (0.55-0.58); PTW: 0.37 (0.35-0.39); WL: 2.102 (2.03-2.22); HFL: 1.44 (1.36-1.53); CI: 1.1049 (1.0325-1.1579); SI: 1.019 (0.9545-1.0472); PI: 1.2719 (1.1538-1.3611); FI: 0.685 (0.6634-0.7073).

**Color**: Head almost entirely red, slightly opaque, with blackening near the anterior mandibular articulation and the frontal carina. The mesosoma and petiolar scale have the same red color as the head. Gaster predominantly black, with slight yellowish tones on the edges of the sternal and tergal plates. Mandibles and antennae are a darker red than the rest of the head, especially in the funiculus area. In most cases, legs show noticeable blackening along their entire length, except for the femur in the area adjacent to the tibiofemoral joint.

**Head**: Slightly taller than wide, subrectangular, with subparallel sides that are slightly broader behind the eyes and the gena converging forward; occipital edges rounded but with a much less pronounced angle than in the major worker. Anterior margin of the clypeus straight or in some cases with a slight notch in the central area, posterior margin bilobed, with a notch in the central section; on the medial sector of the anterior edge of each lobe there are 2 to 3 yellowish setae, and in the more lateral portion a similar number but smaller in size (2 to 3 times smaller) and of a whiter coloration; surface of the clypeal plate micropunctate and with little shine, with small and subtle concavities sparsely scattered across its surface; presence of both long and short setae (3 to 4 times shorter), the former erect and the latter pressed close to the tegument surface (“appressed”), making them difficult to see; presence of a medial keel, which reaches both the anterior and posterior margins of the clypeus, in both cases splitting into two branches very close to the edge towards each lobe of the clypeus. Eyes relatively small in comparison to the size of the head (less so in comparison to the major worker) and slightly oval, approximately 0.7 times as wide as tall, and 0.49 times as tall as the height of the gena. Frontal carina extending from the upper section at the margin of the clypeus to about halfway up the eyes, slightly arched especially in its anterior portion; frontal plate with a thin, superficial central groove; tegument surface similar to that of the clypeus, likewise with long and short setae, and spaced grooves on its surface. Antennal fossa shallow, without special sculpture. Head with tegument similar to that of the clypeus, with long-erect and short-“appressed” setae over the entire surface, including the lateral and posterior margins. Antennal scapes moderately long; almost perfectly straight, approximately as long as the head (SI ∼ 1), with their apex gradually widening; funiculus longer than the scape; pedicel almost the same size as the first flagellar segment. Scape with a very fine micro-reticulation, with lines difficult to perceive, opaque tegument; with small, widely spaced grooves across its surface, similar to the tegument of the head; presence of short, whitish “appressed” pilosity; in frontal view, the lower sector of the condylar bulb of the antenna has abundant very small erect setae, approximately 10 times smaller than the erect setae of the clypeus. Mandibles rounded, with micro-reticulations that increase in size distally, becoming very coarse nearby the teeth; tegument with small, widely spaced grooves; setae of intermediate size relative to the erect setae present on the clypeus, subdecumbent towards the mandibular apex, and small “appressed” setae over the entire surface. **Mesosoma**: Length, 2.1 - 2.47 times longer than wide; dorsum of mesosoma forming a regular arch between the pronotum, mesonotum, and propodeum, without a furrow between them; propodeum without protuberances. Pronotum rounded at its lateral edges; anterior edge flexed towards the dorsal side. Mesosoma in the dorsal sector densely micropunctate and slightly opaque, in the lateral sector the integument is micropunctate and microstriated; anterior edge of the pronotum (neck) microstriated; propodeum micropunctate. Abundant erect setae on the dorsal sector of the pronotum, mesonotum, and propodeum, most abundant on the latter, but not on the sides; “appressed” setae on the entire surface of the pronotum, mesonotum, and propodeum, both dorsally and laterally. **Petiole**: Scale-shaped, thin, 1.52–1.73 times thinner than long; apex rounded throughout its extension; posterior side flat, anterior side slightly convex; presence of 8 long setae, similar in size to those on the clypeus, located on the edge of the petiolar scale, separated into two groups of four by a slight bare recess; microreticulated, slightly opaque integument. **Gaster**: Tegumento con microreticulado muy fino, superficie ligeramente opaca, exceptuando en bordes de placas tergales que presentan mayor brillo; Presencia de setas largas - erectas y cortas - apressed en toda la superficie dorsal y ventral, distribuidas de manera espaciada permitiendo la visualización del tegumento; presencia de línea de setas de tamaño intermedio (¾ a ½ el tamaño de las setas largas) situada a lo largo de todo el borde posterior, cercana de cada placa, tanto en placas tergales como esternales. **Legs**: Posterior femur elongated, 0.66–0.7 times as long as the mesosoma (FI); femur of all legs slightly flattened anteroposteriorly, tibia as well but to a lesser extent; coxa, femur, and tibia covered with long, erect setae, which on the coxa are of the same size and on the femur and tibia are ⅓ or ½ the length of the setae on the clypeus; presence of short pilosity, appressed to the integument, throughout the length of the legs; femur, in the region of the femoro-tibial joint, with four setae: one anterior, one posterior, and two on the dorsal side of the joint, separated from each other, present on all legs. Integument micropunctate and microstriated along the entire length of the legs.

### 2) Camponotus brunnipes sp nov

(Figs. 6-8)

**Fig. 6:**
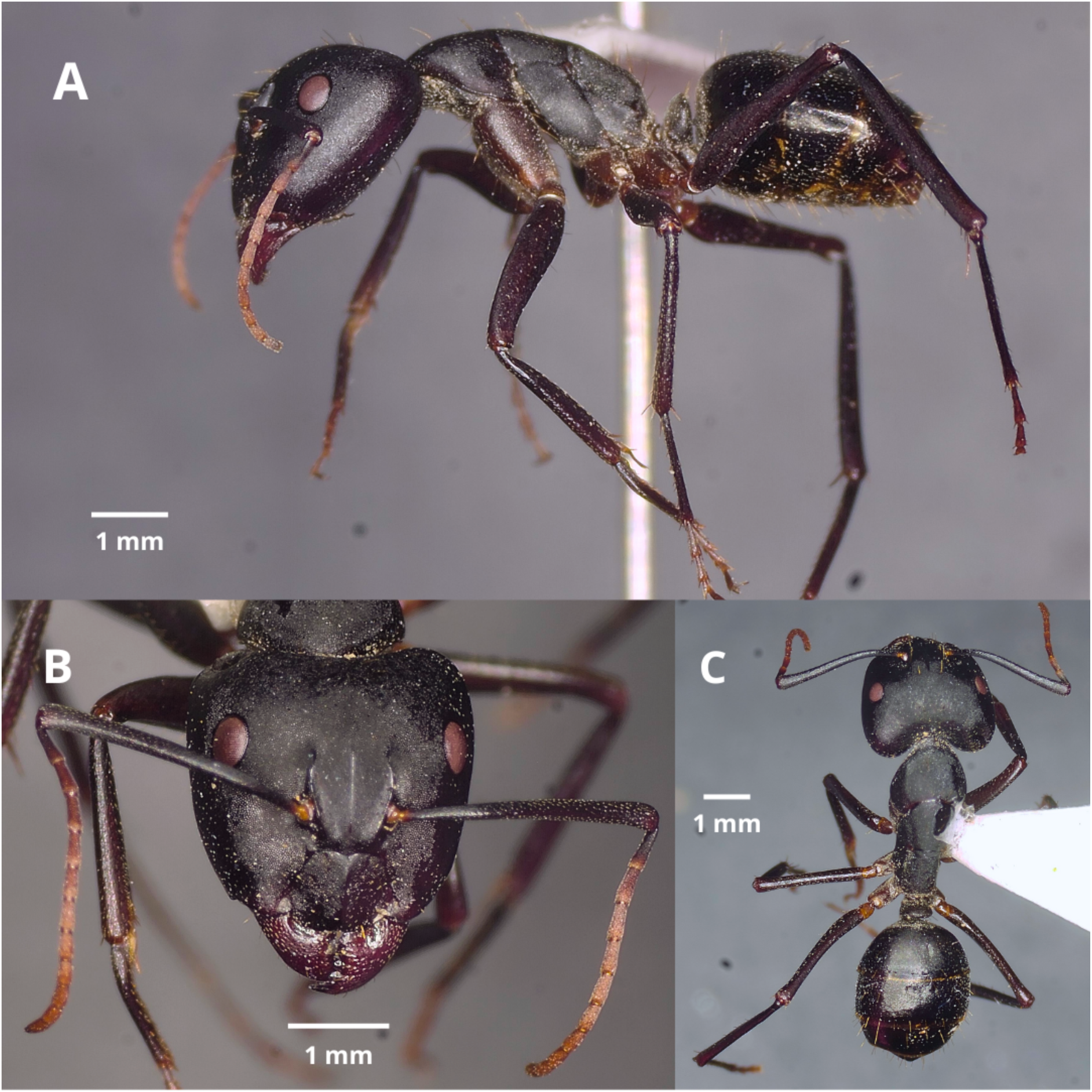
C*a*mponotus *brunnipes* sp nov. major worker photographies (A) Lateral view (B) Frontal / Cephalic view (C) Dorsal view

**Fig. 7:**
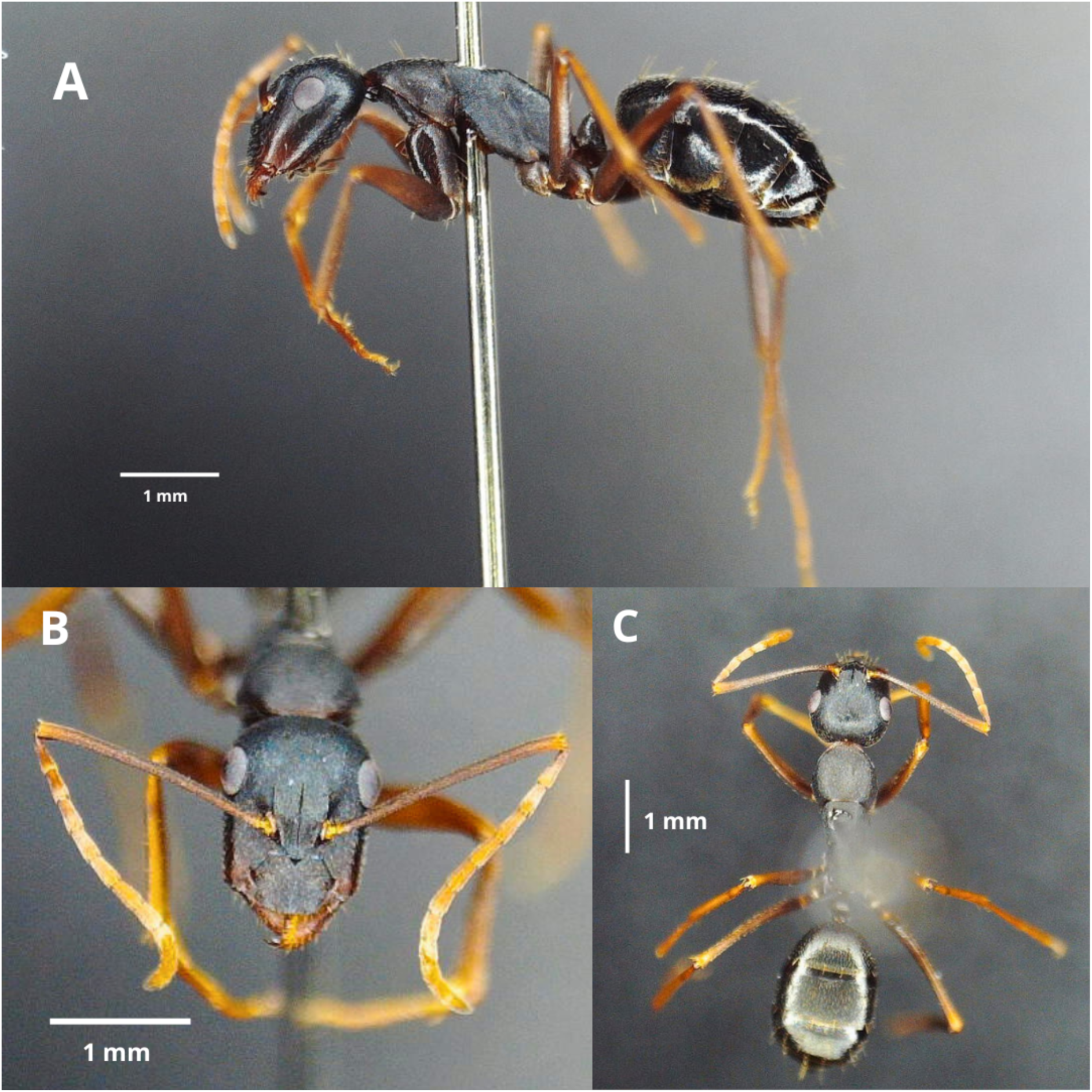
C*a*mponotus *brunnipes* sp nov. minor worker (A) Lateral view (B) Frontal / Cephalic view (C) Dorsal view

**Fig. 8:**
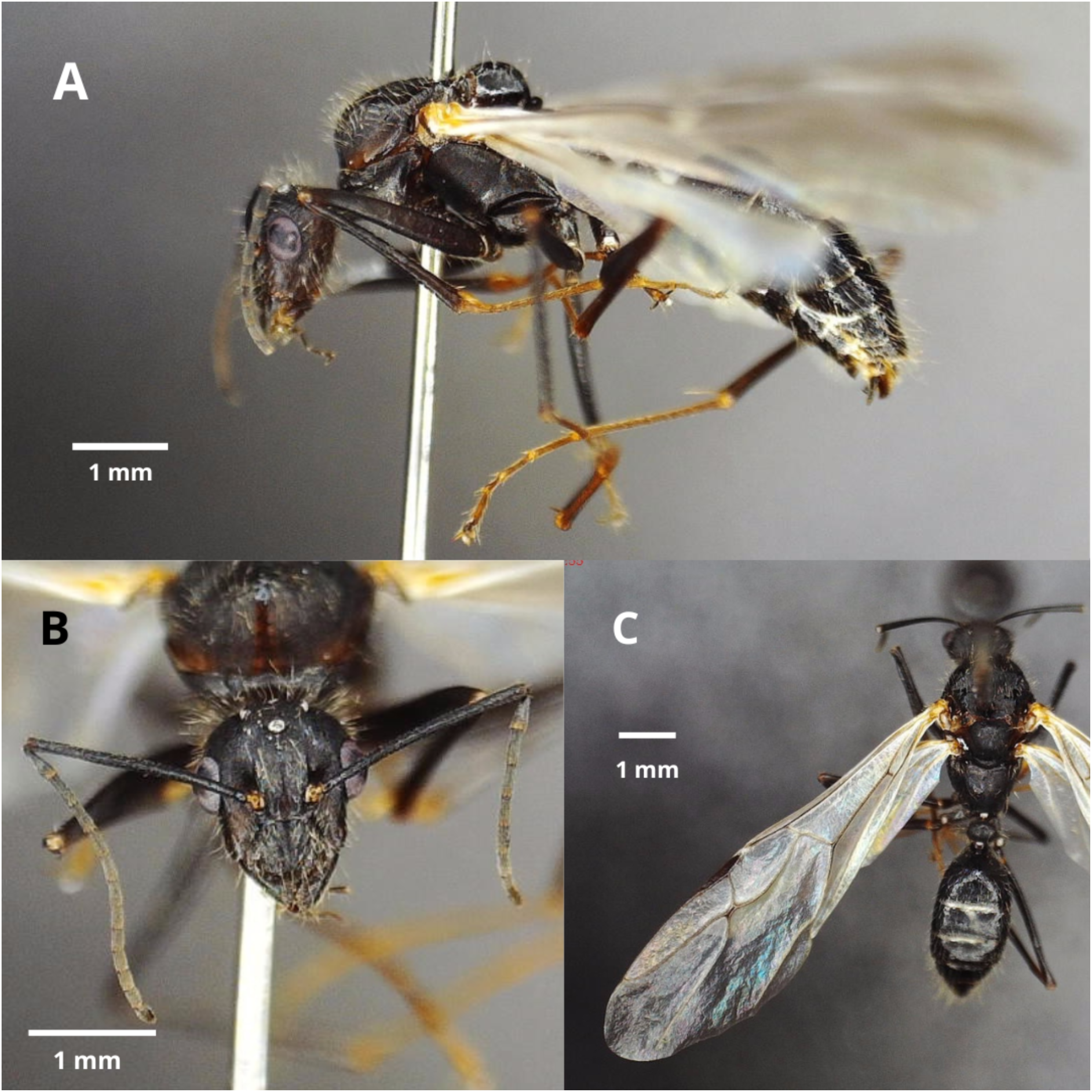
C*a*mponotus *brunnipes* sp nov. male (A) Lateral view (B) Frontal / Cephalic view (C) Dorsal view where the wing venation of the left forewing can be seen.

**Type material (pinned). Holotype:** Major worker (UMCE_IE_1139). Chile, San Clemente (Región del Maule), Cascada Invertida | −35.95473 / −70.57363, 10 IV 2025, B. Arenas (UMCE-IE, UNAP). **3 Paratypes** (UMCE_IE_1140 - UMCE_IE_1144): 2 major workers, 2 minor workers, 1 male: same information as holotype (IE-UMCE).

**Etimology:** Its epithet “brunnipes” refers to the brown color found on the legs of individuals of this species, especially noticeable in minor workers, and on the coxa of major workers.

**Diagnosis:** *Camponotus brunnipes* is a species whose workers are distinguished from the workers of all other species found in Chile by possessing the following set of characteristics: Antennae entirely (in minor workers) or at least the funiculus (in major workers) colored brown/rusty, lighter than the rest of the head. The edges of the gena lack pilosity (setae). Legs are brown, lighter than the head, thorax, and abdomen, almost entirely so in minor workers, and especially marked on the coxae of major workers. The gaster has sparse pilosity (setae), which allows the dark brown/black tegument to remain visible.

#### Description

**Major worker** (n=4, Figs. x-x): **Measurements:** EL: 0.5625 (0.53-0.6); HL: 2.7275 (2.62-2.86); HW: 2.8875 (2.73-3.01); SL: 2.615 (2.48-2.7); PW: 1.6425 (1.58-1.75); PTH: 1.1275 (1.04-1.26); PTW: 0.4125 (0.37-0.45); WL: 3.4775 (3.3-3.8); HFL: 2.7725 (2.63-2.89); CI: 0.945 (0.9169-0.9597); SI: 0.9593 (0.9301-1.0112); PI: 2.7389 (2.5333-2.8919); FI: 0.7987 (0.75-0.8401)

**Color:** Head, mesosoma, and petiolar scale predominantly matte black. Gaster predominantly shiny black, with slight yellowish tones on the edges of the sternal and tergal plates, and long and short setae of a yellowish hue. Mandibles are slightly more reddish than the head, especially near the teeth, but not on the teeth themselves, which are entirely black. Antennal bulb brown/ferruginous in color; scape almost completely black except at its base near the bulb, where it is ferruginous; pedicel black in its most proximal portion, tending toward a more ferruginous color distally; flagellum goes from black-ferruginous to ferruginous from proximal to distal. Legs are dark brown, distinct from the black of the head and mesosoma; the brown coloration is especially lighter in the case of the coxae.

**Head:** Slightly wider than tall, subquadrate, with subparallel sides that are slightly broader behind the eyes and the gena converging forward; occipital margins broadly rounded; posterior margin concave, as a result the posterolateral corners of the head protrude backward. Anterior and posterior margins of the clypeus bilobed, with a notch in the central area; the presence or absence of setae is variable, with no more than one per lobe and/or one emerging from the central notch, relatively short compared to other long setae present on the head; presence of a small notch in the central area of each lobe (bilateral), in some cases with a seta emerging from it; setae are ferruginous in color; integument of the clypeal plate is micropunctate and has little sheen; presence of both long and short setae; in the case of long setae, there are 3 per lobe, arranged in an anteroposterior line: one almost at the posterior margin of the clypeus, another centrally, and another near the anterior margin; in the case of short setae, these are appressed, so small that their visibility is difficult, and they are sparsely and loosely distributed; presence of a median keel, which reaches the anterior margin of the clypeus, extending to the posterior margin where it splits in two towards the edge of each lobe. Eyes are relatively small and oval compared to the size of the head, approximately 0.8 times as wide as tall, and 0.38 times as tall as the height of the gena. Frontal carina extending from the upper area to the margin of the clypeus, reaching approximately halfway up the eyes, slightly arched; frontal plate with a thin, shallow central groove; the surface of the integument similar to the clypeus, likewise with both long and short setae, the former mostly present in the same anteroposterior line as the long setae of the clypeus, extending towards the coronal area of the head; in the coronal area, just above the height of the eyes, there is a slight depression in the integument. Antennal fossa shallow, without special sculpturing. Head with integument similar to that of the clypeus, with short appressed setae over its entire surface, and long erect setae only in the coronal area. Antennal scapes moderately long; in frontal view, almost completely straight, 0.93-1.01 times the width of the head (SI), with the apex gradually widening; funiculus longer than the scape; pedicel elongated relative to the first flagellar segment, scape with very fine micropunctation that is hard to perceive, integument slightly shiny; presence of short, whitish, appressed pilosity; in frontal view, the lower area of the antennal condylar bulb with abundant, very small erect setae, approximately 10 times smaller than the erect setae of the clypeus. Mandibles rounded, with microrreticulations that increase distally, becoming very coarse in the area adjacent to the teeth; integument with small, spaced notches, setae of intermediate size relative to those present in the coronal area of the head, subdecumbent, loosely distributed; and small, appressed setae over their entire surface. **Mesosoma**: Long, 2.09 - 2.17 times longer than wide; dorsum of mesosoma forming a regular arch between the pronotum, mesonotum, and propodeum, without a notch between them; propodeum flat. Pronotum rounded on its lateral edges; anterior edge flexed dorsally. Mesosoma densely micropunctate and slightly opaque; anterior edge of pronotum (neck) microstriated; propodeum micropunctate and microstriated. Numerous erect setae on the dorsal area of the pronotum, mesonotum, and posterior area of the propodeum, but not on the sides; “appressed” setae over the entire surface of the pronotum, mesonotum, and propodeum, both dorsally and laterally. **Petiole**: Con forma de escama, delgado, 1.38-1.7 veces más delgado que ancho; ápice redondeado en toda su extensión; cara posterior plana, cara anterior ligeramente convexa; presencia de 4 setas largas, similares al tamaño de las del clípeo, en el borde de la escama peciolar, separadas en dos grupos de dos por un ligero receso sin pilosidad; tegumento microreticulado ligeramente opaco. **Gaster**: Tegument with very fine micropunctation, surface slightly shiny; Presence of long setae across the entire dorsal and ventral surface, spaced out in such a way that the tegument remains visible; a line of long setae located near the posterior edge of each plate, both on the tergal and sternal plates. **Legs**: Posterior femur elongated, 0.8 times as long as the mesosoma; femur of all legs slightly flattened anteroposteriorly, as are the tibiae but to a lesser extent; coxa, femur, and tibia covered with short “appressed” setae; femur, at the area of the femoro-tibial joint, with four setae present—one anterior, one posterior, and two on the dorsal side of the joint, separated from each other—found on all legs. Tegument micropunctate and microstriated along the entire length of the legs.

**Minor worker** (n=4, Fig 3)

Measurements: EL:0.395 (0.36-0.42); HL: 1.6425 (1.43-1.88); HW: 1.22 (1.03-1.46); SL 1.95 (1.76-2.19); PW: 1.0625 (0.89-1.17); PTH: 0.645 (0.57-0.7); PTW: 0.3175 (0.27-0.37); WL: 2.52 (2.17-2.74); HFL: 2.2075 (1.83-2.45); CI: 1.3517 (1.2877-1.3883); SI: 1.1905 (1.1494-1.2308); PI: 2.0394 (1.8919-2.1212); FI: 0.8742 (0.8433-0.8942)

**Color:** Head, mesosoma, and petiolar scale predominantly matte black, except in some cases where the anterior border of the pronotum (neck) has a brown color. Gaster predominantly shiny black, with slight yellowish tones on the edges of the sternal and tergal plates, and both long and short setae with a yellowish hue. Mandibles are slightly more reddish/brown than the head, especially in the area adjacent to the teeth where the color is notably lighter, but not on the teeth themselves, which are completely black. Antennal bulb light brown, almost yellow in some cases; scape brown, lighter than the head, throughout its length, becoming even lighter closer to its joint with the pedicel; pedicel and flagellum ferruginous in color. Legs are dark brown on the coxae, slightly lighter than the black on the head and mesosoma; trochanter, femur, tibia, and tarsi have a light brown coloration, especially lighter in the case of minor workers.

**Head:** Taller than wide, subrectangular, with almost parallel sides, very slightly wider behind the eyes and with the gena slightly converging towards the front; occipital margins rounded, but with a much less pronounced angle than in the major worker. Posterior clypeal margin bilobed with a notch in the central area, unlike the anterior margin which has a very subtle notch, in some cases almost absent; Presence of 2 to 3 setae per lobe on the anterior edge, in some cases one emerging from the central notch, of intermediate to short length compared to the longer setae present elsewhere on the head; presence of a small notch in the central area of each lobe (bilateral), in some cases with a seta emerging from it; setae with a ferruginous coloration; Clypeal plate tegument micropunctate and with little sheen; presence of both long and short setae; in the case of the long setae, there are 3 per lobe, distributed in an antero-posterior line: one almost at the posterior edge of the clypeus, emerging more medially than the rest, another central, and another almost at the anterior edge; as for the short setae, these are “appressed”, not as small or spaced as in the major worker; presence of a medial keel, which reaches the anterior and posterior edge of the clypeus, where it bifurcates towards the edge of each lobe. Eyes relatively small and oval compared to the size of the head, approximately 0.66 times as wide as high and 0.5 times as high as the height of the gena. Frontal carina extending from the upper area above the posterior edge of the clypeus to about a third of the height of the eyes, slightly arched; frontal plate with a thin and superficial central groove extending up to the same height as the frontal carina; surface of the tegument similar to the clypeus, likewise with the presence of long and short setae, the first mostly present in the same anteroposterior line as the long setae of the clypeus, extending towards the coronal area of the head. Antennal fossa superficial, without special sculpture. Head with tegument similar to that of the clypeus, with short “appressed” setae throughout its surface, and long erect setae only in the coronal area. Antennal scapes long; in frontal view almost completely straight, 1.14– 1.23 times the width of the head (SI), with an almost uniform width along its entire length, slightly wider near the articulation with the pedicel; funiculus longer than the scape; pedicel similar in size to the first flagellar segment, scape with very fine micropunctation that is difficult to perceive, tegument slightly shiny; presence of short whitish “appressed” pubescence; in frontal view, the lower area of the antennal condylar bulb has abundant very small erect setae, approximately 10 times smaller than the erect setae of the clypeus. Mandibles rounded, with micro-reticulations increasing in the distal direction, becoming very rough in the area next to the teeth; tegument with small, spaced notches and setae of intermediate size compared to those found in the coronal area of the head, subdecumbent, distributed in a spaced manner, and small “appressed” setae throughout its surface. **Mesosoma:** Long, 2.31 – 2.44 times longer than wide; dorsum of mesosoma forming a regular arch between pronotum, mesonotum, and propodeum, without indentation between them; propodeum flat. Pronotum rounded on its lateral edges; anterior margin flexed dorsally. Mesosoma densely micropunctate and slightly opaque; anterior margin of pronotum (neck) microstriated; propodeum micropunctate and microstriated. Presence of erect setae on the dorsal section of the pronotum, mesonotum, and in greater density on the propodeum, but not on the lateral sections; “appressed” setae over the entire surface of the pronotum, mesonotum, and propodeum, both dorsally and laterally. **Petiole**: Shaped like a scale, slightly slender, 1.15–1.48 times longer than wide; apex rounded throughout its length; posterior surface flat, anterior surface slightly convex; presence of 4 to 6 long setae, similar in size to those on the clypeus, on the edge of the petiolar scale, separated into two groups by a slight recess without pilosity; microreticulate integument, slightly opaque. **Gaster**: Tegument with very fine micropunctation, slightly shiny surface; presence of long setae over the entire dorsal and ventral surfaces, distributed sparsely, allowing the tegument to be seen; a line of long setae located near the posterior edge of each plate, both on tergal and sternal plates. **Legs**: Elongated posterior femur, 0.75–0.84 times as long as the mesosoma (FI); femur of all legs slightly flattened anteroposteriorly, similarly the tibiae but to a lesser extent; coxa, femur, and tibia covered with short “apressed” setae; femur, at the femoro-tibial joint, with four setae present: one anterior, one posterior, and two on the dorsal side of the joint separated from each other, present on all legs. Tegument micropunctate and microstriated along the entire length of the legs. Tegument micropunctate and microstriated along the entire length of the legs.

### Taxonomic Remarks

Both new species are members of the subgenus *Tanaemyrmex* (Ashmead 1905): High polymorphism, with major workers with broad heads that are usually emarginated towards the posterior end; clypeus with an elongated anterior lobe (Emery 1925). In the territory of Chile all species are considered to be of this subgenus: *Camponotus chilensis* (Spinola 1851)*, Camponotus distinguendus* (Spinola 1851)*, Camponotus morosus* Smith 1858*, Camponotus spinolae* Roger 1863*, Camponotus ovaticeps* (Spinola, 1851) and *Camponotus hellmichi* Menozzi 1935.

## Discussion

The present study substantially increases the number of *Camponotus* species in Chile. Based on the taxonomic history of this group in the country, one might presume that the species richness is low. However, based on the present study describing two new species from the exact same locality, the presumption of low species richness in the country could be challenged. We suggest that the apparently low number of *Camponotus* species present in Chile is likely due, at least in part, to the lack of taxonomists and basic taxonomic work and sampling. We urge the need for increasing government support for taxonomic practice, providing funding and promoting greater opportunities for early career scientists in the field of Systematics and Taxonomy.

## Acknowledgements

The authors thank Mario Elgueta and Maribel Beltrán for their kind assistance, and access to the collections of the Museo Nacional de Historia Natural (MNHN) and the Insect Collection of the Instituto de Entomología, Universidad Metropolitana de Ciencias de la Educación (UMCE-IE), respectively. This work was partially supported by projects DIUMCE-08−2025−ID and FONDECYT (11220706). BA thanks the support from a Beca de Magister ANID.

## Declarations

Conflict of interest. All the authors declare no conflict of interest.

## References

AntWeb. Version 8.112. California Academy of Science, online at https://www.antweb.org. Accessed 10 June 2025

Ashmead, W. H. (1905). A skeleton of a new arrangement of the families, subfamilies, tribes and genera of the ants, or the superfamily Formicoidea. The Canadian Entomologist, 37(11), 381–384.

Bezděčková, K., Bezděčka, P., & Machar, I. (2015). A checklist of the ants (Hymenoptera: Formicidae) of Peru. Zootaxa, 4020(1), 101–133. 10.11646/zootaxa.4020.1.4

Bolton, B. (1995). A New General Catalogue of the Ants of the World. Harvard University Press, Cambridge, M.A.

Braby, M. F., Hsu, Y. F., & Lamas, G. (2024). How to describe a new species in zoology and avoid mistakes. Zoological Journal of the Linnean Society, 20, 1–16. DOI: 10.1093/zoolinnean/zlae043

Emery, C. (1925). Hymenoptera, Fam. Formicidae, subfam. Formicinae. Genera Insectorum, 183.

Fisher, B. L., & Cower, S. P. C. P. (2007). Ants of North America: A Guide to the Genera.

Kusnezov, N. (1951). El género Camponotus en la Argentina (Hymenoptera, Formicidae). Acta Zoológica Lilloana, 12, 183–252.

Mackay, W., & Delsinne, T. (2009). A New Species of Carpenter Ant (Hymenoptera: Formicidae: Camponotus) from Paraguay with a Key to the New World Members of the maculatus Species Complex. Sociobiology, 53(2), 487–498.

Menozzi, C. (1935). Le Formiche del Cile. Zoologische Jahrbücher Abteilung Für Systematik, 67(4), 61 319-336.

Roger, J. (1863). Die neu aufgefülii’ten Gattungen und Arten meines Formiciden-Verzeichnisses. Ergänzung einiger früher gegebenen Beschreibungen, 7, 131–214.

Rohlf, F.J. (2010) tpsDig File Utility Program. Version 2.16. Department of Ecology and Evolution, State University of New York, Stony Brook.

Salata, S., Demetriou, J., Georgiadis, C., & Borowiec, L. (2023). Camponotus Mayr, 1861 (Hymenoptera: Formicidae) of Cyprus: generic synopsis and description of a new species. Asian Myrmecology, 16, 1–33. DOI: 10.20362/am.016007

Smith, F. (1858). Catalogue of the Hymenopterous Insects in the collection of the British Museum. Iv. Formicidae. 216.

Spinola, M. (1851). Formicidae. En: Gay C (ed) Historia física y política de Chile, Zoología. Imprenta de Maulde et Renou, Paris, Francia., 6, 232–246

Wilson EO. 1955. A monographic revision of the ant genus Lasius. Bulletin of the Museum of Comparative Zoology 113:1–201

